# Accurate denoising of single-cell RNA-Seq data using unbiased principal component analysis

**DOI:** 10.1101/655365

**Authors:** Florian Wagner, Dalia Barkley, Itai Yanai

## Abstract

Single-cell RNA-Seq measurements are commonly affected by high levels of technical noise, posing challenges for data analysis and visualization. A diverse array of methods has been proposed to computationally remove noise by sharing information across similar cells or genes, however their respective accuracies have been difficult to establish. Here, we propose a simple denoising strategy based on principal component analysis (PCA). We show that while PCA performed on raw data is biased towards highly expressed genes, this bias can be mitigated with a cell aggregation step, allowing the recovery of denoised expression values for both highly and lowly expressed genes. We benchmark our resulting ENHANCE algorithm and three previously described methods on simulated data that closely mimic real datasets, showing that ENHANCE provides the best overall denoising accuracy, recovering modules of co-expressed genes and cell subpopulations. Implementations of our algorithm are available at https://github.com/yanailab/enhance.

## Introduction

Single-cell RNA-Seq (scRNA-Seq) enables the simultaneous measurement of the transcriptome across thousands of cells from complex tissues and entire organs^1–5^. While unprecedented in its resolution and scale, scRNA-Seq protocols only detect a small fraction of mRNAs in each cell, with estimated detection rates ranging from 3%^6^ to 21%^7^. This limited detection of mRNAs results in high levels of technical noise, referred to as sampling noise^6^. Studies have shown that for all protocols that employ unique molecular identifiers (UMIs) to uniquely identify transcripts, the sampling noise can be accurately approximated using the Poisson distribution, suggesting that the set of transcripts detected from each cell is random^6,8–11^. While sampling noise affects all measurements in a dataset, relative noise levels, quantified using the coefficient of variation (CV), increase in a predictable fashion as the true expression level of a gene decreases^6,10–12^.

Popular methods for visualizing cell similarities in two or three dimensions such as t-SNE^13^ and UMAP^14^ are partly able to overcome sampling noise by defining local neighborhoods of cells and ensuring their proximity in the visualization, effectively aggregating information from similar cells. However, these methods do not generate denoised gene expression values, and thus cannot be used to perform gene-level analyses, for example with scatter plots or heatmaps^15^. To overcome this limitation, a diverse array of methods have been proposed to computationally remove noise by sharing information across cells and genes^10,15–21^. The general lack of an experimental ground truth has made it difficult to establish and compare the accuracies of these approaches, and it is currently unclear what levels of theoretical and algorithmic complexity are necessary or appropriate to address the denoising problem.

Here, we describe *ENHANCE* (**E**xpression de**n**oising **h**euristic using **a**ggregation of **n**eighbors and principal **c**omponent **e**xtraction; https://github.com/yanailab/enhance), a simple and effective algorithm for denoising scRNA-Seq data based on principal component analysis (PCA). We show that PCA on raw scRNA-Seq data is strongly biased towards highly expressed genes, and that this bias can be mitigated by inserting a cell aggregation step. Moreover, we describe a simulation-based approach for efficiently estimating the number of principal components carrying biological information in scRNA-Seq data. After validating our algorithm using CITE-Seq data, we propose a novel approach to simulate scRNA-Seq data with realistic technical and biological characteristics, and use it to systematically benchmark ENHANCE and three previously described denoising methods.

## Results

To develop a strategy for denoising scRNA-Seq datasets containing a complex mixture of cell populations with unknown degrees of similarity, we observed that while true expression differences between cell types and subpopulations result in substantial gene-gene correlation structure^22,23^, sampling noise affects the expression measurement of each gene in a largely independent fashion. We therefore reasoned that principal component analysis (PCA), applied to variance-stabilized data^10^, could be a powerful approach for separating biological heterogeneity from technical noise in scRNA-Seq data. By inferring expression levels based upon the leading principal components (PCs), we would retain only true biological differences, while discarding the technical noise captured by higher PCs (**Fig. 1a**). However, we encountered two problems with this approach. First, since technical noise becomes an increasingly dominant factor as the true expression level of a gene decreases, the signal captured by PCA is strongly biased towards highly expressed genes (**Fig. 1b**). Second, the number of PCs that capture biological differences can vary between datasets, and it is unclear how to determine this number in a principled and efficient manner. To address the first problem, we begin with a nearest-neighbor aggregation step, in which the expression profile of each cell is replaced by the aggregate sum of profiles of itself and the *k-1* most similar other cells (neighbors), where *k* is chosen in relation to the average transcript count and the number of cells in the dataset (**Methods**). Aggregation reduces the noise levels, particularly for lowly expressed genes, and therefore results in a substantial mitigation of the PCA expression bias (**Fig. 1b**). To address the second problem, we developed a simulation approach to estimate the maximum amount of technical noise that a single PC can capture. We then only retain “significant” PCs that capture at least twice this amount, to ensure that most of their signal represents biological differences (**Fig 1c**). Our resulting denoising algorithm (**Fig. 1a** and **Methods**) does not require any parameter tuning and executes in under two minutes on the datasets examined in this study.

**Figure 1.**
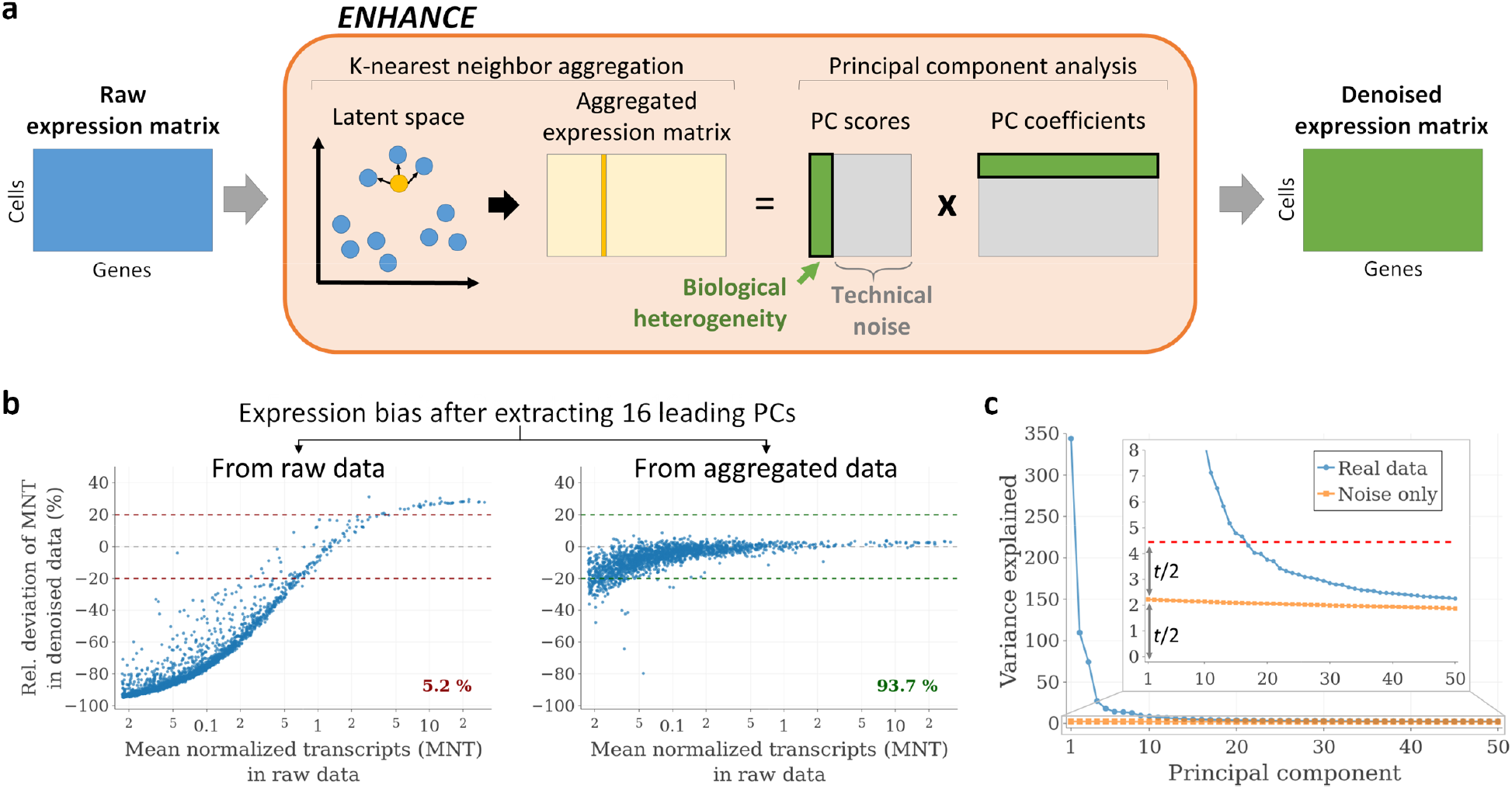
Outline and of the ENHANCE denoising algorithm. **a** Schematic overview of the ENHANCE algorithm. **b** Per-gene expression bias after extracting the leading PCs from a PBMC dataset (4,334 cells), before and after K-nearest neighbor aggregation, with *k*=58. The percent of expressed genes with an absolute bias of 20% or less (relative to the raw data) is indicated. For better readability, only a random subset of 2,000 genes is shown. **c** Variance explained for the first 50 principal components (PCs), for the same PBMC dataset as well as for simulated data containing only technical noise. The threshold (*t*; red dotted line) for determining significant PCs is twice the amount of variance explained by the first PC of the simulated data.

To validate our approach, we decided to take advantage of a CITE-Seq dataset of human peripheral blood mononuclear cells (PBMCs), which comprises single-cell expression measurements for the transcriptome as well as a panel of cell surface proteins. We applied ENHANCE to the single-cell mRNA measurements and observed a dramatic effect on the expression profile of individual genes such as the naïve T cell marker *CCR7* (**Fig. 2a**). To determine whether the denoised expression profiles more accurately represented the true expression levels than the raw profiles, we used the protein expression data to independently define naïve and memory subsets of T cells (**Fig 2b, c**). Using these subsets, we found that unlike in the raw data, *CCR7* expression levels in the denoised data predicted naïve T cell identity with high sensitivity and specificity. We compared the ENHANCE results for *CCR7* to those of MAGIC^15^, SAVER^16^, and ALRA^21^, and found that only ALRA provided a comparable improvement (**Fig. 2d** and **Methods**). We next expanded our analysis to all genes with an AUROC > 0.6 in the raw data, and found that on average, ENHANCE led to the best AUROC improvement, followed by MAGIC and SAVER (**Fig. 2e**). The success of ENHANCE depended on both nearest-neighbor aggregation and PC extraction, as omitting either step resulted in smaller increases in AUROC scores (**Supplementary Fig. 2**). Denoising provided the most benefit for genes with small absolute expression differences between the cell types, underscoring the dramatic impact of noise on the T cell expression profiles (**Fig. 2f**).

**Figure 2:**
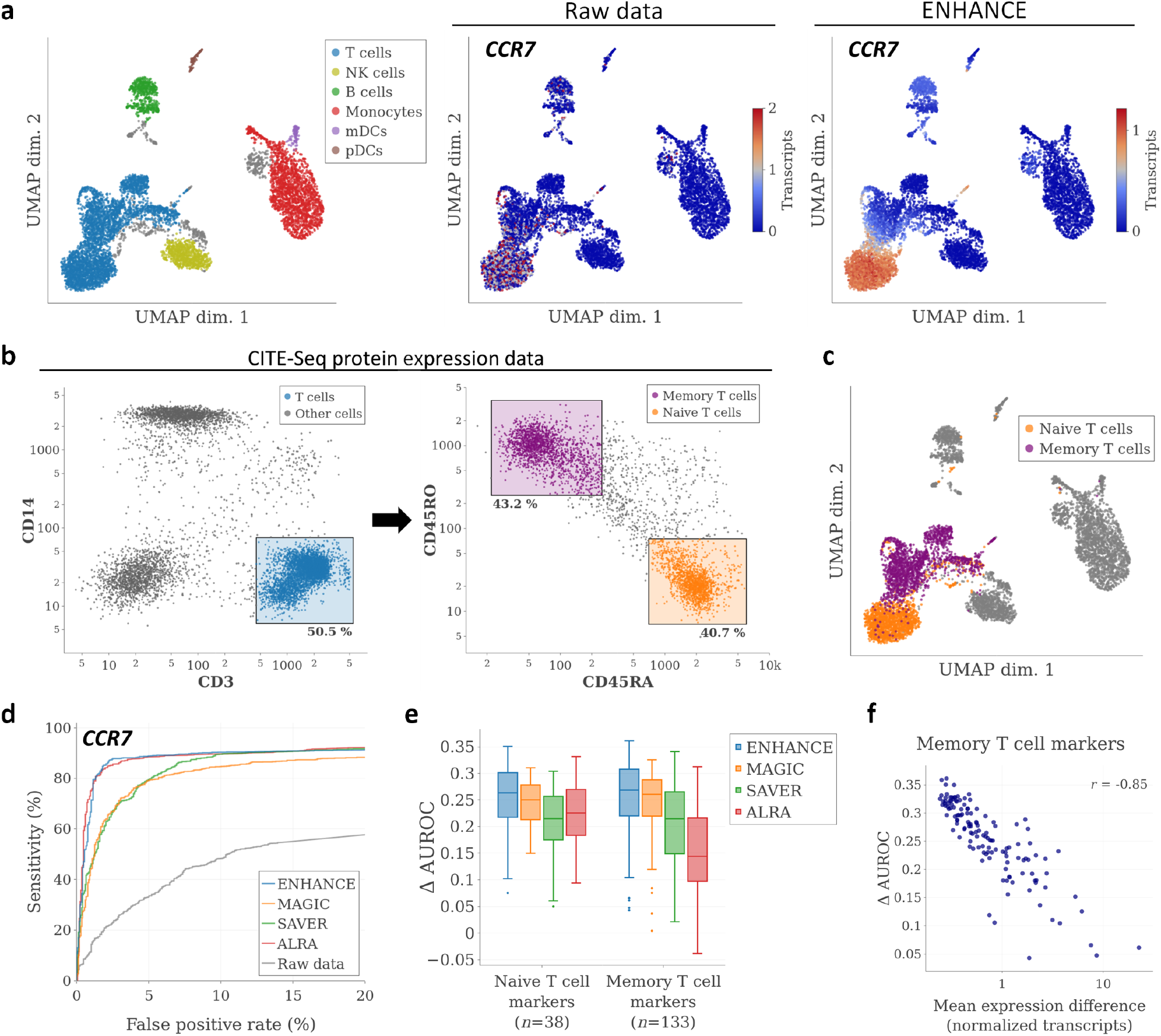
Validation of ENHANCE using CITE-Seq data. **a** Left: UMAP visualization and cell type assignments for transcriptome data from a CITE-Seq PBMC dataset (7,666 cells). Unassigned cells are shown in gray. Middle and right: Expression profile of *CCR7* in the raw data and after denoising with ENHANCE. To improve readability, the color scale for raw *CCR7* expression values was clipped at 2. **b** “Gating” strategy for identifying naïve and memory T cell populations in the same dataset based on protein expression data. **c** Gating results overlaid on top of the UMAP visualization shown in (a). **d** ROC curves showing performance of CCR7 expression as a marker for distinguishing between naïve and memory T cells. **e** Improvement in AUROC scores for ENHANCE and three other denoising methods, for all genes with AUROC > 0.6 in the raw data. **f** Correlation between improvement in AUROC score and absolute expression difference between naïve and memory T cells.

We next aimed to systematically compare the accuracies of ENHANCE, MAGIC, SAVER, and ALRA in a simulation study. To ensure that the results were representative of the methods’ performance on real scRNA-Seq data, our goal was to generate artificial datasets whose biological and technical sources of variation mirrored those of real datasets. To overcome the limitations of previously described simulation approaches^15,16,18–20,24^ (**Supplementary Note 1**), we used the output of a denoising method as the ground truth, and then simulated realistic efficiency and sampling noise (**Methods**). We applied this approach to simulate data based on a renal cell carcinoma biopsy sample^25^, which contained a heterogeneous set of populations from the tumor microenvironment (**Fig. 3a**). To avoid biasing the analysis towards any single denoising method, we performed three separate simulations, Sim-ENHANCE, Sim-MAGIC, and Sim-SAVER, based on the outputs of ENHANCE, MAGIC and SAVER, respectively. To test whether each simulated dataset captured the biological differences present in the real dataset, we performed clustering on the real dataset and overlaid the cluster assignments onto an independent t-SNE visualization of the simulated dataset. We observed a nearly perfect congruence between clusters in the real and the simulated datasets (**Fig. 3a**). We next assessed the technical characteristics of the simulated data by comparing the means, standard deviations, and fraction of zero values for each gene with those in the real data, and again found very close agreements (**Fig. 3b**). Thus, the simulated datasets recapitulated the cell type differences present in the real dataset, while simultaneously exhibiting the characteristic noise profile of scRNA-Seq data. At the same time, the simulated datasets were far from a carbon copy of the real data, as between 68-70% of the total variance in the simulated datasets constituted randomly generated noise (**Methods**).

**Figure 3.**
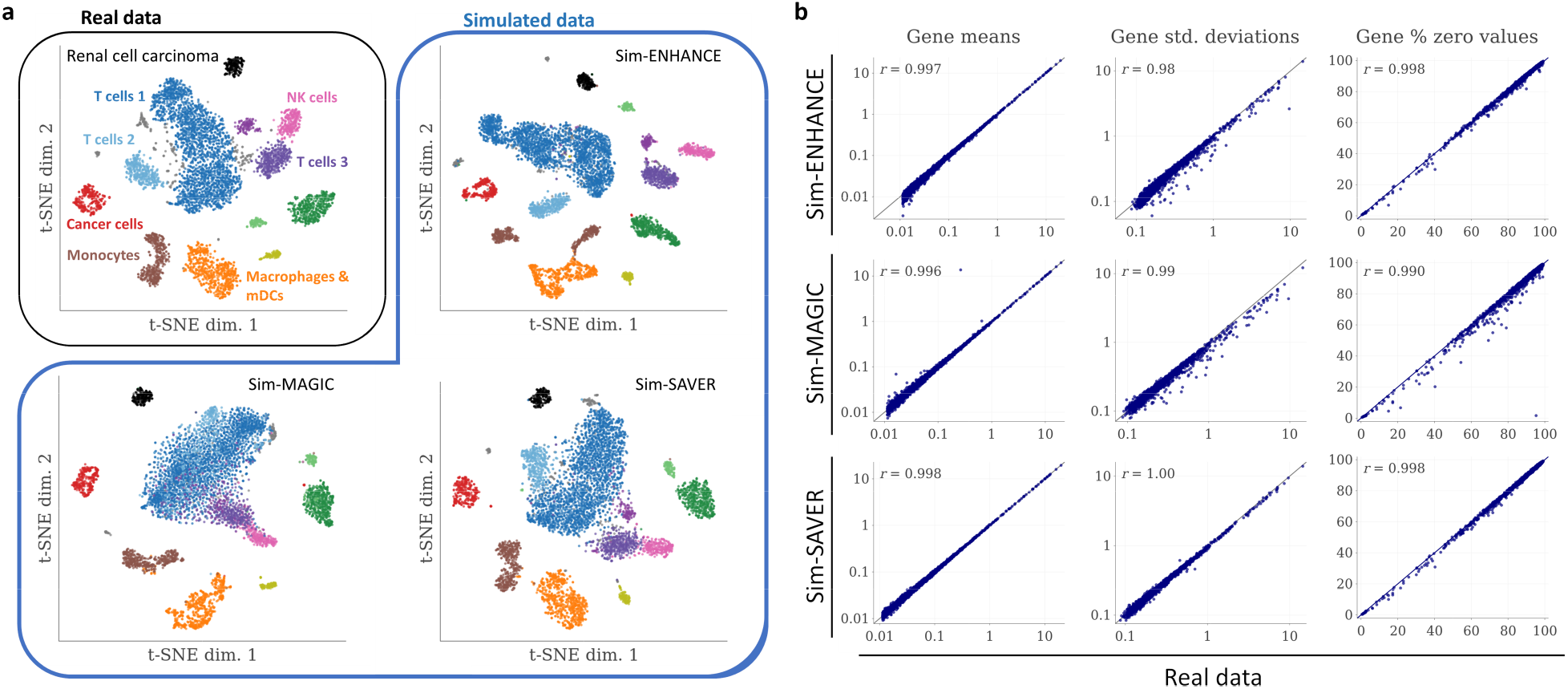
Simulation of realistic scRNA-Seq data based on a renal cell carcinoma dataset. **a** Top left: t-SNE visualization and clustering results for a renal cell carcinoma dataset (6,232 cells). Rest: Independent t-SNE visualizations of data simulated with Sim-ENHANCE, Sim-MAGIC and Sim-SAVER, respectively, overlaid with the clustering results from the real data. **b** Comparison of gene means, standard deviations and proportion of zero values between real and simulated data.

After confirming that our simulated datasets had realistic biological and technical properties, we next aimed to compare the accuracies of ENHANCE, MAGIC, SAVER, and ALRA. To this end, we first used the ground truth data from each simulation to identify variable genes (**Methods**), and then assessed the correlation between their expression patterns in the denoised data and the ground truth. We grouped genes by expression level, and found that as expected, denoising accuracies were generally the lowest for lowly expressed genes (**Fig. 4a**). In both Sim-ENHANCE and Sim-MAGIC, SAVER and ALRA were clearly outperformed by ENHANCE and MAGIC, which exhibited similar performance. In Sim-SAVER, SAVER outperformed the other methods on highly expressed genes, but not on genes with intermediate and low expression levels. We observed that SAVER generally failed to accurately denoise many lowly expressed or less variable genes (**Supplementary Fig. 3**). Next, we aimed to examine the ability of each method to recover expression differences between closely related cell types. To this end, we took advantage of the fact that the kidney cancer dataset contained at least three different populations of T cells. We therefore selected only the T cells from each dataset, and recalculated the correlation between ground truth and denoised gene expression patterns. The results showed that ENHANCE outperformed MAGIC in both Sim-ENHANCE and Sim-MAGIC, on genes with high and intermediate expression (**Fig. 4a**). We also used heatmaps to examine the expression patterns of genes with high and variable expression in T cells, which provided a more direct view of the differences in accuracy (**Fig. 4b**). In particular, the heatmaps showed instances where MAGIC resulted in a substantial loss of subpopulation differences (“over-smoothing”; red box in **Fig. 4b**). In contrast, for SAVER and ALRA, smooth expression patterns in the ground truth sometimes exhibited significant variability in the denoised data, suggesting that these methods failed to fully denoise the simulated raw data (“under-smoothing”; blue boxes in **Fig. 4b**). We found again that the accuracy of ENHANCE depended on both cell aggregation and PC extraction (**Supplementary Fig. 4**). To assess the robustness of our findings, we repeated our analyses on technical replicates, and obtained nearly identical results (**Supplementary Fig. 5**). Additionally, we performed a second benchmark study using simulated PBMC data, and obtained similar results (**Supplementary Fig. 6**). In summary, our analyses demonstrated that ENHANCE exhibited the best overall denoising accuracy, and that it specifically outperformed the other methods in recovering expression differences between closely related cell types.

**Figure 4.**
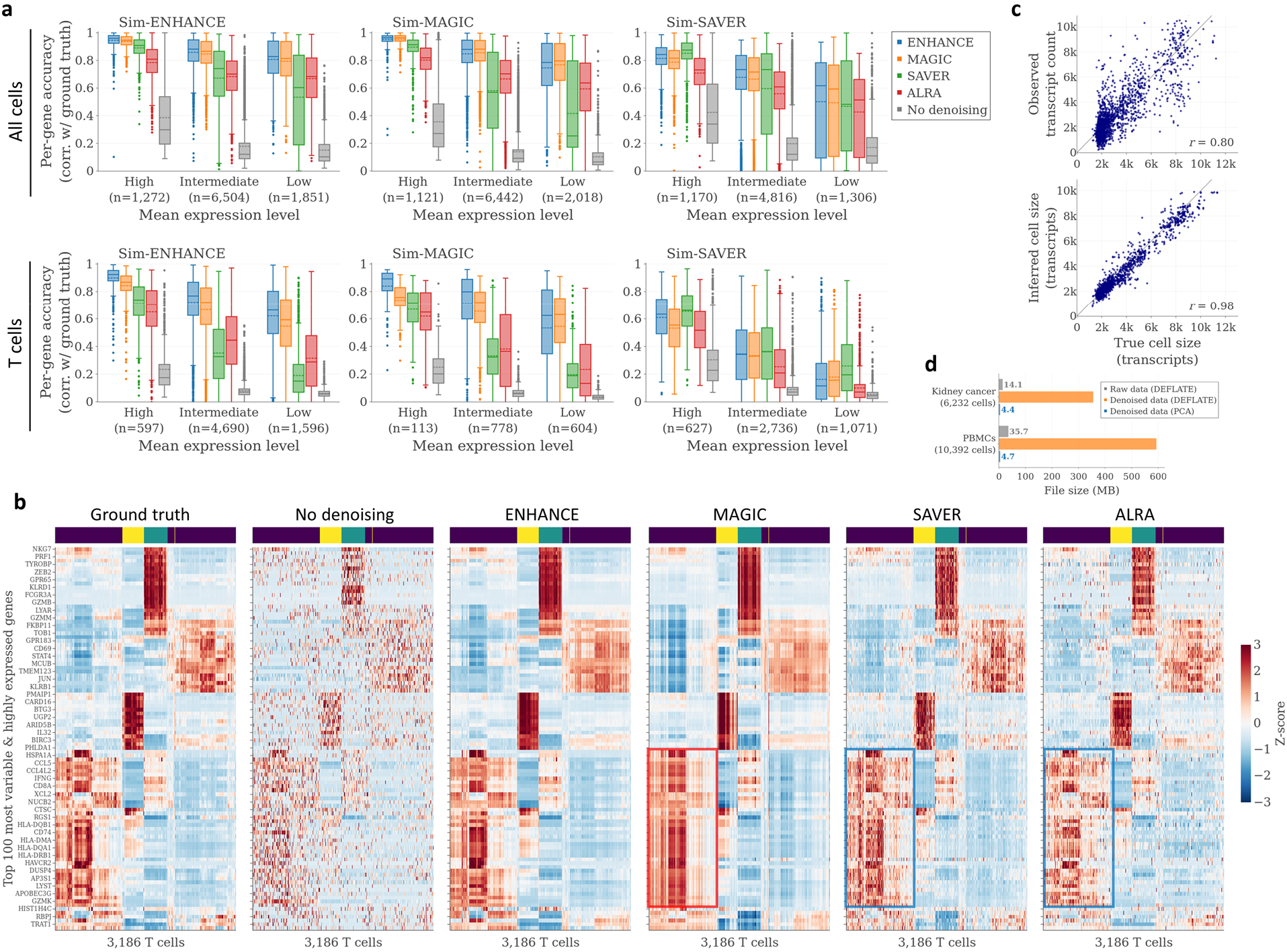
Benchmarking of ENHANCE and three other denoising methods on simulated renal cell carcinoma data. **a** Box plots showing gene-wise correlations between ground truth and denoised data, for simulated renal cell carcinoma data generated using three different simulation methods. Genes are grouped by expression level, and only variable genes are included in the analysis (**Methods**). Top: Correlations calculated using all cells. Bottom: Correlations calculated using only T cells. **b** Heatmaps showing expression patterns for 100 genes across all 3,186 T cells for the Sim-ENHANCE ground truth, the simulated raw counts, and the denoised datasets. All datasets were first normalized and then the expression values for each gene were transformed to z-scores. Above each heatmap, the color-coded T cell cluster identity from **Fig. 2a** is shown for each cell (navy blue=T cells 1, turquoise=T cells 2, yellow=T cells 3). The 100 most variable genes from the “highly expressed” gene category are shown in the heatmap, and example are shown, and genes and cells were ordered according to hierarchical clustering of the ground truth data (with correlation distance and average linkage). **c** Comparison of true, observed, and ENHANCE-inferred cell sizes for the simulated kidney cancer dataset generated using Sim-ENHANCE (each dot is a cell; only a random subset of 2,000 cells is shown). **d** Comparison of file sizes for different ways storing the raw and denoised data in compressed form.

While the other denoising methods only generate normalized expression profiles, ENHANCE also aims to remove efficiency noise and infer differences in cell size. In Sim-ENHANCE, we used bootstrapping in conjunction with the observed cell sizes to simulate efficiency noise. Using this approach, we found that ENHANCE was able to accurately infer cell sizes (**Fig. 4c**). Finally, we noted that since the denoised datasets were no longer sparse, they were not amenable to efficient compression and required more than ten times more disk space than the raw data. However, as the output of ENHANCE can be represented as the scaled product of two much smaller matrices containing PC coefficients and scores, we found that it was possible to reduce the amount of required disk space by about two orders of magnitude, thus allowing for the convenient storage and exchange of ENHANCE results (**Fig. 4d**).

## Discussion

We have described ENHANCE, an effective algorithm for denoising heterogeneous scRNA-Seq datasets that consists of a series of simple steps and relies heavily upon PCA to separate biological signals from technical noise. As an initial validation of our method, we have used CITE-Seq data to assess the extent to which the denoising improves the differential expression of genes between T cell subsets. We then introduced a new approach for simulating scRNA-Seq data that allows real scRNA-Seq datasets to be used as “templates” for generating artificial datasets exhibiting realistic technical noise and biological cell type differences. Using this approach, we performed a benchmark study of our algorithm and three previously described methods, which showed that ENHANCE had the best overall denoising accuracy.

While it remains challenging to accurately recover expression differences between closely related cell types, particularly for lowly expressed genes, our work demonstrates that the denoising problem can be effectively addressed with standard methods such as PCA, and that an increase in theoretical or algorithmic complexity does not automatically lead to more accurate results, mirroring similar findings for the problem of cell type classification^26^. Indeed, low-complexity methods may provide clearer opportunities for optimization. For example, in the case of ENHANCE, a less conservative criterion for selecting the number of significant PCs may prove beneficial in more accurately capturing differences between similar cell types or denoising less abundant cell populations.

The inherent noisiness of scRNA-Seq data has served as a major motivation and a key assumption in the development of many specialized methods for visualization, clustering, and trajectory inference^27–30^. Here, we have demonstrated the ability to accurately denoise scRNA-Seq data without prior analysis steps. This begs the question of whether current scRNA-Seq analysis workflows can be revised to more effectively explore gene expression heterogeneity. For example, hierarchically clustered expression heatmaps^31^ have been a cornerstone in the analysis of bulk gene expression data, but their effectiveness is severely limited when high levels of noise are present. Following denoising, the ability to identify modules of co-expressed genes and cell subpopulations using heatmaps is greatly improved. Additionally, our work quantitatively demonstrates how PCA can be used to capture most of the biological heterogeneity in scRNA-Seq data in an unbiased fashion, thus providing a new foundation for machine learning tasks that rely on low-dimensional data representations.

## Methods

A detailed description of the methods is provided in the **Online Methods**. Python and R implementations of ENHANCE can be found at https://github.com/yanailab/enhance and https://github.com/yanailab/enhance-R, respectively.

## Supporting information

Manuscript with Online Methods

