## Supplementary material for "Accurate denoising of single-cell RNA-Seq data using unbiased principal component analysis": Online Methods

**Principal component analysis.** All principal component analyses were performed on normalized and variance-stabilized data. For normalization, we scale the expression profile of each cell so that the total sum of transcript counts for all cells equals the same number C, which we set to the median total transcript count of the dataset. We therefore refer to this procedure as "median normalization". For variance stabilization, we apply the Freeman-Tukey transform, $y = \sqrt{x} + \sqrt{x+1}$, which has been shown to provide good results for Poisson-distributed data[1,2]. Given the number of leading principal components of interest *d* (see below for our approach for determining *d*), we then perform a truncated PCA using a randomized SVD algorithm[3] as implemented in the *decomposition.PCA* class from the Python package *scikit-learn*, version 0.20[4]. We therefore initialize the *PCA* object with *n_components*=*d* and *svd_solver*='randomized'. The results of the PCA comprise the 1) PC scores $\boldsymbol{S}$, an *n*-by-*d* matrix with coordinates of each cell in PC space, 2) the components $\boldsymbol{W}$, a *d*-by-*p* matrix that contains the coefficients that define each PC, and 3) the gene means $\boldsymbol{\mu}^{trans}$ that were subtracted before determining the singular values and vectors.

**Determining the number of significant principal components.** To determine the number of "significant" principal components for given dataset with an *n*-by-*p* expression matrix $\boldsymbol{X}$, we perform PCA on a simulated "noise expression matrix" $\boldsymbol{X}^{noise}$ with the same dimensions as the real dataset. In this noise matrix, the only source of variance is sampling noise, and there exist no true biological differences between the cells. The idea is to use this noise matrix to derive a PC variance threshold $t^{noise}$, and to determine the number *d* of significant PCs of the real dataset as the number of PCs that explain at least $t^{noise}$ units of variance. (Since PCs are ordered by the amounts of variance explained, this always corresponds to the *d* leading PCs).

We generate $\boldsymbol{X}^{noise}$ in the following way. First, we apply median normalization to the real expression matrix (see above). For each gene, we then calculate the mean transcript count across all cells. We use the resulting mean expression profile $\boldsymbol{\mu}$ as the basis for our simulation. To generate a single simulated expression profile, we simulate the transcript count of each gene by drawing from a Poisson distribution with parameter $\lambda=\mu_i$, where represents the *i*'th element of $\boldsymbol{\mu}$. We repeat this procedure *n* times to generate all *n* expression profiles of $\boldsymbol{X}^{noise}$.

To obtain $t^{noise}$ from $\boldsymbol{X}^{noise}$, we first choose an upper bound for the number of PCs that we would be interested in. We refer to this number as *max_components*. For our analyses, we use *max_components*=50, as we did not expect the datasets analyzed here to have more than 50 meaningful PCs. We then perform PCA with *d*=*max_components* on $\boldsymbol{X}^{noise}$ as described above, with the exception that we are normalizing to the median total transcript count of the real expression matrix. We then set $t^{noise}$ to twice the amount of variance explained by the first principal component. Finally, we perform PCA with *d*=*max_components* on $\boldsymbol{X}$, and determine the number of significant PCs as the number of PCs that capture at least $t^{noise}$ units of variance. Our reasoning for setting $t^{noise}$ to twice the amount of variance captured by the first PC of the noise matrix is that this should guarantee that at least 50% of the variance explained by each "significant" PC of the real data represents true biological differences, rather than technical noise. In other words, each PC of the real data can overfit and capture some technical noise, but we generally would not expect this level to be larger than $t^{noise}$ / 2. Therefore, we generally expect the inclusion of those PCs in the analysis to improve (rather than worsen) the biological fidelity of any analyses based on the PCA results.

**Aggregation of nearest neighbors.** To overcome expression bias when inferring true gene expression levels from principal components, we replace the expression profile of each cell with an aggregated expression profile that combines the UMI counts of the cell itself as well as its k-1 nearest neighbors. As the measurements are independent, this results in a theoretical reduction of noise levels (CV) by a factor of $\sqrt{k}$. To find an appropriate value of *k* for a given dataset, we make use of our observation that a median total transcript count of 200,000 transcripts (for the aggregated expression matrix) leads to sufficiently unbiased PCA results (**Fig. 1b**). We therefore define $k^{TC} = \lceil 200{,}000/\ C \rceil$, where *C* is the median total transcript count of the raw expression matrix. To avoid choosing a k that would result in "oversmoothing", i.e., the aggregation of cells from different subpopulations, we do not allow k to exceed a certain fraction of the total number of cells in a dataset, which we refer to as $f^{max}$. We therefore use $k = \min\left(k^{TC}, \lfloor f^{max} * n \rfloor\right)$, where *n* is the number of cells in the dataset. We use $f^{max} = 2\%$, however this value can be adjusted according to the known or expected abundances of cell populations in the data.

Given a raw expression matrix and a matrix containing PC scores for all cells, we identify the *k* nearest neighbors of each cell (including the cell itself) based on Euclidean distance calculated using the PC scores. We then calculate an aggregated expression profile for each cell by summing up the transcript counts in the raw data of its *k* nearest neighbors in a gene-wise fashion. We refer to the *n*-by-*p* matrix containing all aggregated expression profiles as the "aggregated expression matrix". We also calculate an "estimated cell size" for each cell, by calculating the median total transcript count of all of its neighbors. By using the median, this estimate is robust to cases in which a minority of the identified neighbors belong to a different cell population with a potentially different size.

**Extraction of principal components.** Based on the results of a PCA with *d* leading principal components (see above), we extract the transcript counts represented by these *d* principal components in the following way. First, we multiply the PC scores $\boldsymbol{S}$ with the components $\boldsymbol{W}$, again obtaining an *n*-by-*d* matrix $\boldsymbol{T}^{center}$. We then add back the gene means $\boldsymbol{\mu}^{trans}$, resulting in the matrix **T** that represents normalized and Freeman-Tukey-transformed (FT-transformed) transcript counts. To reverse the Freeman-Tukey transformation, we then set all values of $\boldsymbol{T}$ that are below 1 to 1, since values below 1 represent technical artifacts that do not correspond to valid FT-transformed transcript counts. We then apply the inverse of the Freeman-Tukey transform, $y = (x^2 - 1)^2/(4x^2)$, to each element of $\boldsymbol{T}$. This results in a matrix $\boldsymbol{D}^{norm}$ that represents the normalized transcript counts as captured by the first *d* PCs.

**ENHANCE algorithm.** The ENHANCE algorithm can be conceptually divided into three phases. In the first phase, the number of significant PCs is determined (*num_components*). In the second phase, an aggregated expression matrix is generated. In the third phase, the signal captured by the first *num_components* PCs of the aggregated matrix is extracted, and then the resulting expression profiles are scaled to match the estimated cell sizes. The first phase (determining the number of significant PCs) is fully described above, and we will therefore skip its description here. The inferred number of significant principal components is used for all PCAs performed in phases two and three.

The goal of the second phase is the generation of an aggregated expression matrix, which then allows for application of PCA in a way that is not strongly biased towards the most highly expressed genes. To improve the accuracy with which neighbors are identified in the aggregation step (see above), we realized that the PCs obtained by applying PCA on the raw data will inevitably exhibit overfitting, i.e., capture some portion of technical noise, especially for the higher PCs. To reduce the amount of overfitting and thus improve the accuracy with which correct neighbors are identified when closely related subpopulations are present, we perform two PCA/aggregation cycles. The second PCA is performed on the results of the first aggregation step (after scaling back to the median transcript count of the raw data), and the raw data are then projected onto the resulting PCs. The resulting PC scores are then used in the second aggregation step, which again operates on the raw data.

In the third and final phase, a PCA is conducted on the aggregated expression matrix, and the signal corresponding to the significant PCs is extracted, resulting in an $n$-by-$p$ matrix $\boldsymbol{D}^{norm}$, as described above. Finally, the resulting expression profile of each cell is scaled so that its total transcript count matches its estimated cell size, which was calculated during the second aggregation step (see above). This produces the final denoised expression matrix $\boldsymbol{D}$.

**Single-cell RNA-Seq preprocessing workflow.** To ensure consistent preprocessing of all scRNA-Seq datasets analyzed in this study, we aimed to apply a uniform scRNA-Seq preprocessing workflow to all datasets. The goal of this workflow is to only retain known protein-coding genes that are located on the nuclear genome (not the mitochondrial genome), and to remove cells that do not meet certain quality control criteria. First, we extracted a list of known protein-coding genes from the human Ensembl genome annotations (e.g., release 92; http://ftp.ensembl.org/pub/release-92/gtf/homo_sapiens/Homo_sapiens.GRCh38.92.gtf.gz). We then curated a list of 13 protein-coding genes located on the mitochondrial genome as all genes that have a name starting with "MT-". The first step of our preprocessing workflow consists of removing all genes that are not contained in our list of protein-coding genes (we identify genes using their Ensembl IDs). In a second step, we remove all cells with a total transcript count of less than 1,000 (counting only transcripts from the protein-coding genes). We also remove cells where the 13 mitochondrially encoded genes account for more than 15% of the observed transcripts. This removes cells with low-quality data, represented by a low total transcript count and/or a high proportion of mitochondrially encoded transcripts, potentially due to cell lysis.

**Quantification of gene expression bias after extracting leading principal components.** To quantify the expression bias after extracting the leading PCs (**Fig. 1b**), we relied on a PBMC dataset labeled "4k PBMCs from a Healthy Donor", which belonged to a series of datasets made available by 10X Genomics under the heading "Chromium Demonstration (v2 Chemistry)" (https://support.10xgenomics.com/single-cell-gene-expression/datasets). We downloaded the UMI-filtered expression matrix from the 10x Genomics website (http://cf.10xgenomics.com/samples/cell-exp/2.1.0/pbmc4k/pbmc4k_filtered_gene_bc_matrices.tar.gz), and applied our scRNA-Seq preprocessing workflow (see above; using genome annotations from Ensembl release 92), which resulted in a dataset containing 4,334 cells with a median transcript count of 3478.5. We then applied the ENHANCE algorithm, as well as a modified version in which we skipped the kNN-Aggregation step (by manually setting $k$=1). To quantify expression bias, we first normalized both the raw data as well as the outputs of the ENHANCE algorithm to the median total transcript count of the cells in the raw data. We then calculated the ratios between the gene means in the normalized ENHANCE output data and the gene means in the normalized raw data. We also calculated a mean expression threshold to determine which genes we considered "expressed". To calculate this threshold $t$, we multiplied the median total transcript count with 5% and 100 TPM ($t = C*0.05*0.001$). Thus, a gene will generally be considered expressed if it is expressed at a concentration of at least 100 TPM in at least 5% of the cells in the data. This resulted in 8,861 expressed genes for the PBMC dataset.

**Selection and application of other denoising algorithms.** We compared ENHANCE to MAGIC[5], SAVER[6], and ALRA[7]. We chose MAGIC and SAVER as they enjoy high impact and popularity based on their publication in high-profile journals, high number of citations, and high number of "stars" on their GitHub repositories. We included ALRA as it is included in the highly popular Seurat package[8], making it easily accessible to many researchers in the field, and because it also adopts a PCA-based approach to denoising. We used the following packages and workflows. SAVER was performed using version 1.1.1 of the SAVER package, and with the parameter "estimates.only=TRUE". ALRA was performed using version 3.0 of the Seurat package, by applying the function "Seurat:::RunALRA" to a normalized Seurat object. To obtain denoised expression values on a linear scale, we applied the function y=exp(x) - 1 to the results, thus inverting the log-transformation with pseudocount 1 that is part of Seurat's normalization procedure. Both SAVER and the SEURAT packages were obtained from CRAN, and run under Microsoft R Open 3.5. MAGIC was performed using the Python MAGIC package v1.5.5, obtained from GitHub (https://github.com/KrishnaswamyLab/MAGIC/, commit 84c052d). Following a warning that unexpressed genes may have a negative impact on the results, we first removed all unexpressed genes from each expression matrix. We then followed the steps described in the EMT tutorial provided by the authors on GitHub. Briefly, we applied the function "scprep.normalize.library_size_normalize" to the matrix. We did not apply a square root transformation to the normalized data, as we found that this resulted in much worse denoising accuracies. We note that the tutorial is not clear on whether to perform this step or not. A comment indicates that this step should be included, however the results in the notebook show results obtained using untransformed data. Finally, we noticed that the denoised matrices contained some negative values. As negative expression levels interfere with standard normalization procedures and have no meaningful interpretation, we set those values to zero. Finally, we added the unexpressed genes back to the matrix, with all zero values.

**Validation of denoising performance using CITE-Seq PBMC data.** To validate the ENHANCE algorithm, we relied on a PBMC CITE-Seq dataset labeled "10k PBMCs from a Healthy Donor - Gene Expression and Cell Surface Protein", which belonged to a series of datasets made available by 10X Genomics under the heading "Chromium Demonstration (v3 Chemistry)" (https://support.10xgenomics.com/single-cell-gene-expression/datasets). We downloaded the UMI-filtered expression matrix and separated the transcriptome from the protein measurements, thus producing a transcriptome and a protein expression matrix (the measurements for the 17 proteins were easily identified as their gene identifiers were the only ones that did not start with "ENSG…"). We then applied our scRNA-Seq preprocessing workflow to the transcriptome expression matrix (see above; using genome annotations from Ensembl release 95), which resulted in a dataset containing 7,666 cells with a median transcript count of 3277.5. We then filtered the protein expression matrix to only retain the measurements from these cells.

To visualize the transcriptome expression data, we first applied PCA with $d$=30 (see above). We then applied UMAP on the PC scores (Python *umap-learn* package version 0.3.8, https://pypi.org/project/umap-learn/). We used the default UMAP parameters (*num_neighbors*=15; Euclidean distance metric), except for setting *min_dist* to 0.5 (default: 0.1). To obtain cell type clusters, we first performed t-SNE on the PC scores (as implemented in the Python *scikit-learn* package, version 0.20.3) with default parameters (*perplexity*=30). We then applied DBSCAN (*scikit-learn*) with *eps*=5.21

and *min_samples*=77 and manually labeled the resulting clusters by examining the expression patterns of marker genes. Specifically, the markers we relied on were *CD3D* (T cells), *KLRD1* (*CD94*; NK cells), *CD79B* (B cells), *CD14* (monocytes), *CD74* (mDCs), and *JCHAIN* (pDCs).

To define naïve and memory T cell populations based on the protein expression data, we first applied median normalization to the protein expression matrix, and then defined T cells are cells with a normalized CD3 UMI count between 450 and 5500, and a normalized CD14 UMI count between 6 and 75 (as shown in **Fig. 1e**, left). This resulted in the identification of 3,870 T cells. We then filtered the raw protein expression matrix for these T cells, and re-applied median normalization. We then defined naïve T cells as those with a normalized CD45RA UMI count between 700 and 6000, and a normalized CD45RO count between 7 and 75. Conversely, we defined memory T cells as those with a normalized CD45RA UMI count between 22 and 250, and a normalized CD45RO count between 250 and 3,500.

To determine the extent to which CCR7 expression discriminates between naïve and memory T cells in the raw data, we selected the cells defined as naïve and memory T cells from the raw transcriptome data, applied median normalization, and generated a ROC curve using the *metrics.roc_curve* function from scikit-learn. To test the ability of each denoising method to remove technical noise while preserving biological expression differences, we repeated this procedure on the denoised data, obtained by applying denoising to the *entire* transcriptome data, i.e. the dataset including *all* cells, not only those defined as naïve or memory T cells. (The results of these analyses are shown in **Fig. 1g**). To systematically examine the improvement of marker-based discrimination between naïve and memory T cells (**Fig. 1h**), we first used the raw expression profiles of the naïve and memory T cells, and applied the *metrics.roc_auc_score* function from scikit-learn[4] v0.20.3 to calculate the area under the ROC curve (AUROC) for either naïve or memory T cells. For each cell type, we then selected the top 10 genes with the highest AUROC scores, and repeated the calculation of AUROC scores using the denoised data (again obtained by applying ENHANCE to the entire dataset).

**Simulation of single-cell RNA-Seq data.** To simulate realistic scRNA-Seq data, we decided to use real scRNA-Seq datasets as a template by applying either ENHANCE, MAGIC, or SAVER and treating the resulting denoised matrix as the ground truth, and then simulating a noisy "raw" gene expression matrix with the same number of cells as the real dataset. We refer to these simulation methods as Sim-ENHANCE, Sim-MAGIC, and Sim-SAVER, respectively. To simulated efficiency noise, we adopted two different approaches. As MAGIC and SAVER do not infer cell sizes, we simulated efficiency noise in Sim-MAGIC and Sim-SAVER by scaling the expression profile of each cell in the ground truth to the total transcript count of the same cell in the real dataset. ENHANCE also infers the true size of each cell. In Sim-ENHANCE, we first calculated a scale factor for each cell, by dividing the cell's total transcript count in the real data by its total transcript count in the denoised data. We then obtained efficiency factors for each cell in the simulated dataset by applying bootstrapping (sampling with replacement) to this set of scale factors. We then scaled the expression profile of each cell in the ground truth by its simulated efficiency factor. After scaling the ground truth expression profiles to simulate efficiency noise, we used a common approach to simulate sampling noise in all three simulation methods. To obtain the simulated transcript count for the *j*'th gene in the *i*'th cell of the simulated matrix, we sampled from a Poisson distribution with $\lambda=e_{ij}$, where $e_{ij}$ represents the value of the *j*'th gene in the *i*'th ground truth expression profile (after scaling).

To quantify the proportion of total variance constituting technical noise in the simulated data, we first median-normalized and FT-transformed the ground truth, calculated the variance for each gene, and calculated the sum total ("true variance"). We then normalized the simulated data to the median total transcript count of the ground truth, applied the FT-transform, calculated the variance for each gene, and calculated the sum total ("simulated variance"). We then subtracted the true variance from the simulated variance, and divided by the simulated variance.

**Benchmarking of denoising accuracies on simulated data.** To benchmark the accuracy of ENHANCE, MAGIC, SAVER, and ALRA, we obtained single-cell RNA-Seq data (generated using the 10X Chromium v2 protocol), from a study of by Young et al. of human kidney cancer[9]. Specifically, we used data from renal cell carcinoma patient 2, tumor sample 2, which consisted of two scRNA-Seq libraries (labeled "RCC2_Kid_T_ldc_2_1" and "RCC2_Kid_T_ldc_2_2" in the annotation file provided by the authors), which we combined and processed using our preprocessing workflow, using Ensembl v92 genome annotations (see above). This resulted in a dataset containing 6,232 cells and 16,532 protein-coding genes, with a median transcript count of 2,251.5.

We then applied the three simulation methods described above to generate three simulated datasets. To assess the accuracy of each denoising method, we then applied each denoising method to each of the simulated datasets. We next aimed to assess the correlation between expression patterns in the denoised data and the ground truth. As the measure of correlation is only meaningful for genes that exhibit a variable expression pattern, we decided to identify variable genes as follows. First, we identified expression genes as those genes with a mean expression value of 0.05 or higher in the ground truth, where the mean is calculated after normalizing the expression matrix to 10,000 transcripts per cell. We reasoned that this represented a very low threshold for expression, as it can be satisfied when a gene has an expression level of 100 TPM in only 5% of the cells in the data. Using this threshold, the number of expressed genes for each of the three simulation methods was approximately 10,000. Next, we further selected variable genes as those genes with a coefficient of variation (CV) of at least 20%. We again reasoned that this represented a permissive threshold, and we found neither denoising method performed well on genes with even lower variabilities. For these variable genes, we then calculated the Pearson correlation between their expression patterns in the ground truth and the denoised matrix.

To evaluate the accuracy of each denoising method separately on genes with high, intermediate, and low expression levels, we defined lowly expressed genes as those with a mean expression of less than 0.1 transcripts (calculated after normalizing the matrix to 10,000 transcripts per cell), genes with intermediate expression level as those with a mean of at least 0.1 but less than 1 transcripts, and highly expressed genes as those with a mean of at least 1 transcript per cell. The results of this analysis are shown in **Fig. 2c**. To evaluate the ability of each denoising method to correctly infer expression differences among closely related cell types, we repeated this evaluation after selecting the three T cell populations that we identified in our clustering analysis (**Supplementary Fig. 1**). The results of this analysis are shown in **Fig. 2d**.

To assess the robustness of our results, we repeated our analysis for technical replicates of our simulated datasets, obtained by repeated the simulation with a different random number generator seed (**Supplemental Fig. 7**). The technical replicates therefore contain the same biological heterogeneity, but independent technical noise (except for the efficiency noise that was modeled using the total transcript counts from the real data). We also repeated the analysis on data simulated based on an entirely different dataset, namely the dataset labeled “10k PBMCs from a Healthy Donor (v3 chemistry)”, provided online by 10X Genomics and belonging to the same series of datasets as the previously described PBMC CITE-Seq dataset. After preprocessing using Ensembl rel. 95 genome annotations, this dataset contained 10,392 cells and 16,334 protein-coding genes, with a median transcript count of 5,831. We evaluated denoising accuracies in an identical way as before, and the results are shown in **Supplemental Fig. 8**.

To evaluate the accuracy with which ENHANCE was able to infer the true size of each cell, we took advantage of the fact that in Sim-ENHANCE, we explicitly simulated efficiency noise. We therefore compared the true, simulated, and inferred cell sizes using the Pearson correlation coefficient (**Fig. 2e**).

**Comparison of data compression efficiencies for raw and denoised data.** To assess how much disk space can be saved when taking advantage of the fact that the ENHANCE result can be represented a the scaled produced of two small matrices containing PC coefficients and scores, respectively, we first had to establish a baseline. While raw scRNA-Seq data is often stored in plain-text form using the MatrixMarket format for sparse matrices, following compression with the DEFLATE algorithm[10], denoised data does no longer exhibit a high degree of sparsity, and a sparse data format would therefore not be an economical choice for storing denoised data. However, we found that an excellent compression ratio for raw scRNA-Seq data can also be achieved by storing the expression matrix in binary form, again followed by the compression with DEFLATE. We reasoned that this approach is equally valid for storing denoised data, and would therefore provide a good baseline for our comparison. We therefore compared the size of a binary file containing the PCA results and the inferred cell sizes (represented as 32-bit floating point values and stored using numpy’s *savez* function), to DEFLATE-compressed binary files containing either the raw expression matrix (represented as 32-bit unsigned integers) or the denoised expression matrix (represented as 32-bit floating point values), in both cases stored using numpy’s *savez_compressed* function.. The results for the kidney cancer and PBMC datasets evaluated in the simulation study are shown in **Fig. 2f**.
