## Supplementary material for "Accurate denoising of single-cell RNA-Seq data using unbiased principal component analysis": Manuscript with Online Methods

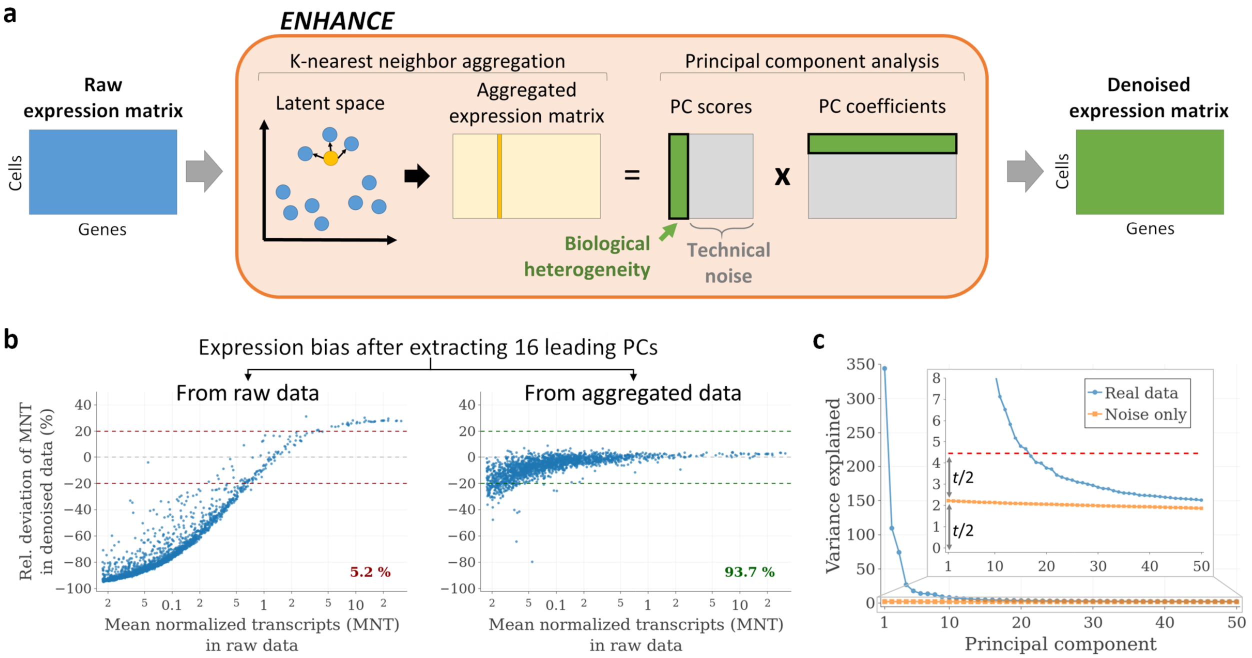


**Figure 1. Outline and of the ENHANCE denoising algorithm. a** Schematic overview of the ENHANCE algorithm. **b** Per-gene expression bias after extracting the leading PCs from a PBMC dataset (4,334 cells), before and after K-nearest neighbor aggregation, with *k*=58. The percent of expressed genes with an absolute bias of 20% or less (relative to the raw data) is indicated. For better readability, only a random subset of 2,000 genes is shown. **c** Variance explained for the first 50 principal components (PCs), for the same PBMC dataset as well as for simulated data containing only technical noise. The threshold (*t*; red dotted line) for determining significant PCs is twice the amount of variance explained by the first PC of the simulated data.

To validate our approach, we decided to take advantage of a CITE-Seq dataset of human peripheral blood mononuclear cells (PBMCs), which comprises single-cell expression measurements for the transcriptome as well as a panel of cell surface proteins. We applied ENHANCE to the single-cell mRNA measurements and observed a dramatic effect on the expression profile of individual genes such as the naïve T cell marker *CCR7* (**Fig. 2a**). To determine whether the denoised expression profiles more accurately represented the true expression levels than the raw profiles, we used the protein expression data to independently define naïve and memory subsets of T cells (**Fig 2b, c**). Using these subsets, we found that unlike in the raw data, *CCR7* expression levels in the denoised data predicted naïve T cell identity with high sensitivity and specificity. We compared the ENHANCE results for *CCR7* to those of MAGIC[15], SAVER[16], and ALRA[21], and found that only ALRA provided a comparable improvement (**Fig. 2d** and **Methods**). We next expanded our analysis to all genes with an AUROC > 0.6 in the raw data, and found that on average, ENHANCE led to the best AUROC improvement, followed by MAGIC and SAVER (**Fig. 2e**). The success of ENHANCE depended on both nearest-neighbor aggregation and PC extraction, as omitting either step resulted in smaller increases in AUROC scores (**Supplementary Fig. 2**). Denoising provided the most benefit for genes with small absolute expression differences between the cell types, underscoring the dramatic impact of noise on the T cell expression profiles (**Fig. 2f**).

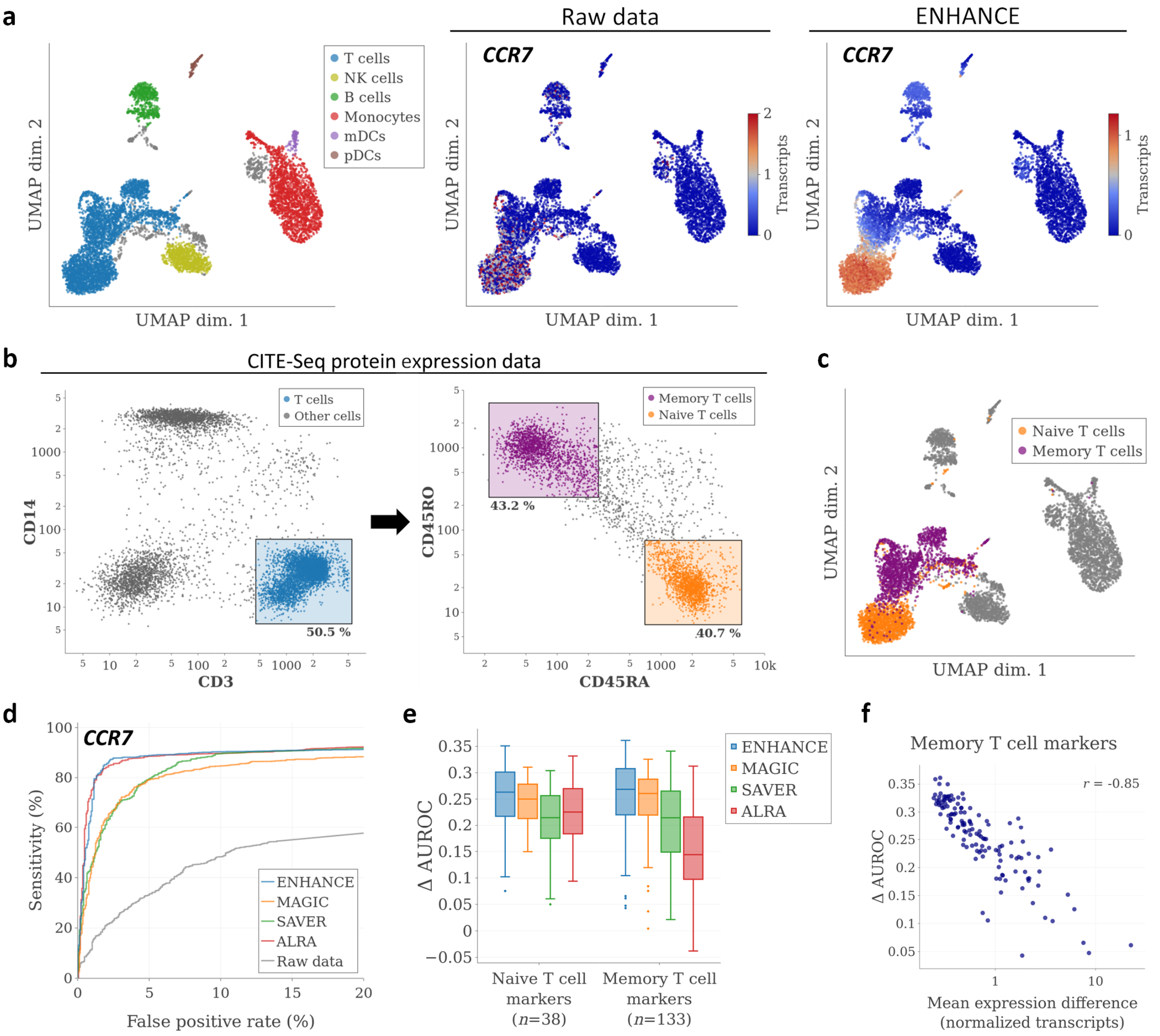


**Figure 2: Validation of ENHANCE using CITE-Seq data. a** Left: UMAP visualization and cell type assignments for transcriptome data from a CITE-Seq PBMC dataset (7,666 cells). Unassigned cells are shown in gray. Middle and right: Expression profile of *CCR7* in the raw data and after denoising with ENHANCE. To improve readability, the color scale for raw *CCR7* expression values was clipped at 2. **b** "Gating" strategy for identifying naïve and memory T cell populations in the same dataset based on protein expression data. **c** Gating results overlaid on top of the UMAP visualization shown in (a). **d** ROC curves showing performance of CCR7 expression as a marker for distinguishing between naïve and memory T cells. **e** Improvement in AUROC scores for ENHANCE and three other denoising methods, for all genes with AUROC > 0.6 in the raw data. **f** Correlation between improvement in AUROC score and absolute expression difference between naïve and memory T cells.

We next aimed to systematically compare the accuracies of ENHANCE, MAGIC, SAVER, and ALRA in a simulation study. To ensure that the results were representative of the methods' performance on real scRNA-Seq data, our goal was to generate artificial datasets whose biological and technical sources of variation mirrored those of real datasets. To overcome the limitations of previously described simulation approaches[15,16,18–20,24] (**Supplementary Note 1**), we used the output of a denoising method as the ground truth, and then simulated realistic efficiency and sampling noise (**Methods**). We applied this approach to simulate data based on a renal cell carcinoma biopsy sample[25], which contained a heterogeneous set of populations from the tumor microenvironment (**Fig. 3a**) . To avoid biasing the analysis towards any single denoising method, we performed three separate simulations, Sim-ENHANCE, Sim-MAGIC, and Sim-SAVER, based on the outputs of ENHANCE, MAGIC and SAVER, respectively. To test whether each simulated dataset captured the biological differences present in the real dataset, we performed clustering on the real dataset and overlaid the cluster assignments onto an independent t-SNE visualization of the simulated dataset. We observed a nearly perfect congruence between clusters in the real and the simulated datasets (**Fig. 3a**). We next assessed the technical characteristics of the simulated data by comparing the means, standard deviations, and fraction of zero values for each gene with those in the real data, and again found very close agreements (**Fig. 3b**). Thus, the simulated datasets recapitulated the cell type differences present in the real dataset, while simultaneously exhibiting the characteristic noise profile of scRNA-Seq data. At the same time, the simulated datasets were far from a carbon copy of the real data, as between 68-70% of the total variance in the simulated datasets constituted randomly generated noise (**Methods**).

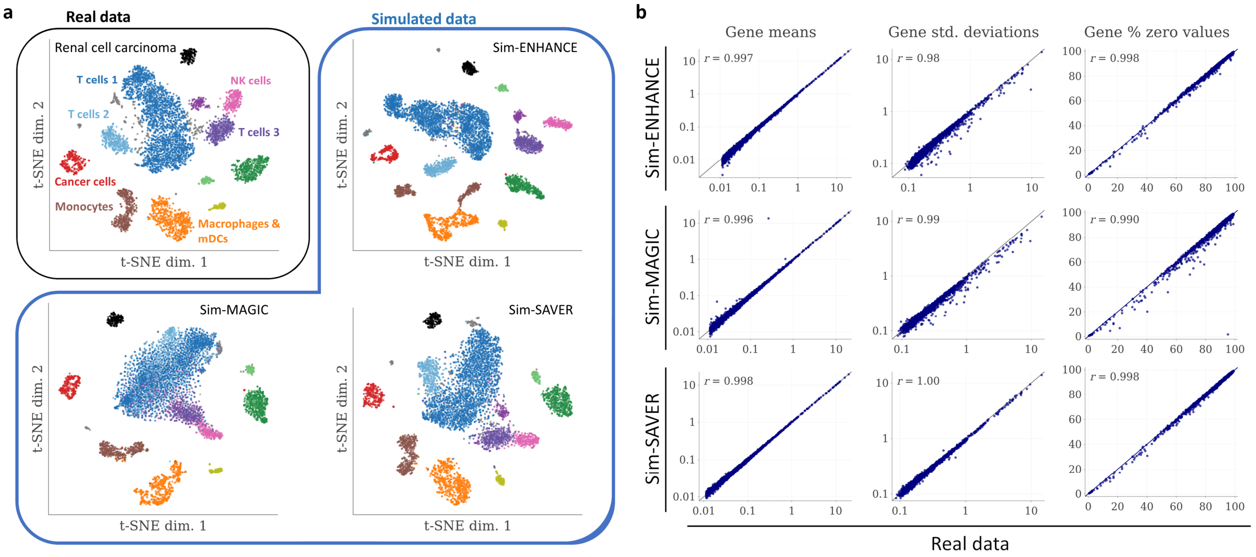


**Figure 3. Simulation of realistic scRNA-Seq data based on a renal cell carcinoma dataset. a** Top left: t-SNE visualization and clustering results for a renal cell carcinoma dataset (6,232 cells). Rest: Independent t-SNE visualizations of data simulated with Sim-ENHANCE, Sim-MAGIC and Sim-SAVER, respectively, overlaid with the clustering results from the real data. **b** Comparison of gene means, standard deviations and proportion of zero values between real and simulated data.

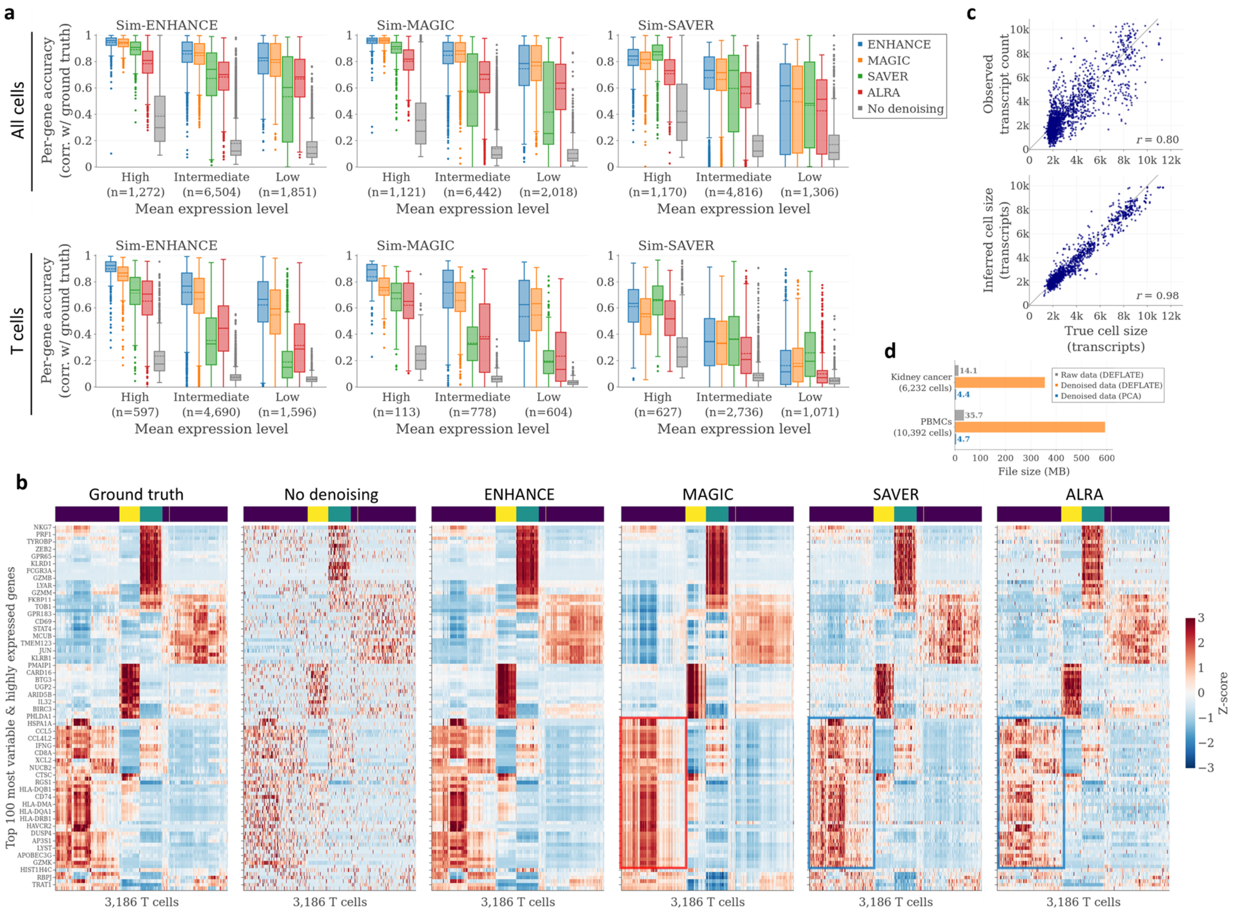


**Figure 4. Benchmarking of ENHANCE and three other denoising methods on simulated renal cell carcinoma data.** **a** Box plots showing gene-wise correlations between ground truth and denoised data, for simulated renal cell carcinoma data generated using three different simulation methods. Genes are grouped by expression level, and only variable genes are included in the analysis (**Methods**). Top: Correlations calculated using all cells. Bottom: Correlations calculated using only T cells. **b** Heatmaps showing expression patterns for 100 genes across all 3,186 T cells for the Sim-ENHANCE ground truth, the simulated raw counts, and the denoised datasets. All datasets were first normalized and then the expression values for each gene were transformed to z-scores. Above each heatmap, the color-coded T cell cluster identity from **Fig. 2a** is shown for each cell (navy blue=T cells 1, turquoise=T cells 2, yellow=T cells 3). The 100 most variable genes from the "highly expressed" gene category are shown in the heatmap, and example are shown, and genes and cells were ordered according to hierarchical clustering of the ground truth data (with correlation distance and average linkage). **c** Comparison of true, observed, and ENHANCE-inferred cell sizes for the simulated kidney cancer dataset generated using Sim-ENHANCE (each dot is a cell; only a random subset of 2,000 cells is shown). **d** Comparison of file sizes for different ways storing the raw and denoised data in compressed form.
