## Supplementary Figures for "Accurate denoising of single-cell RNA-Seq data using unbiased principal component analysis"

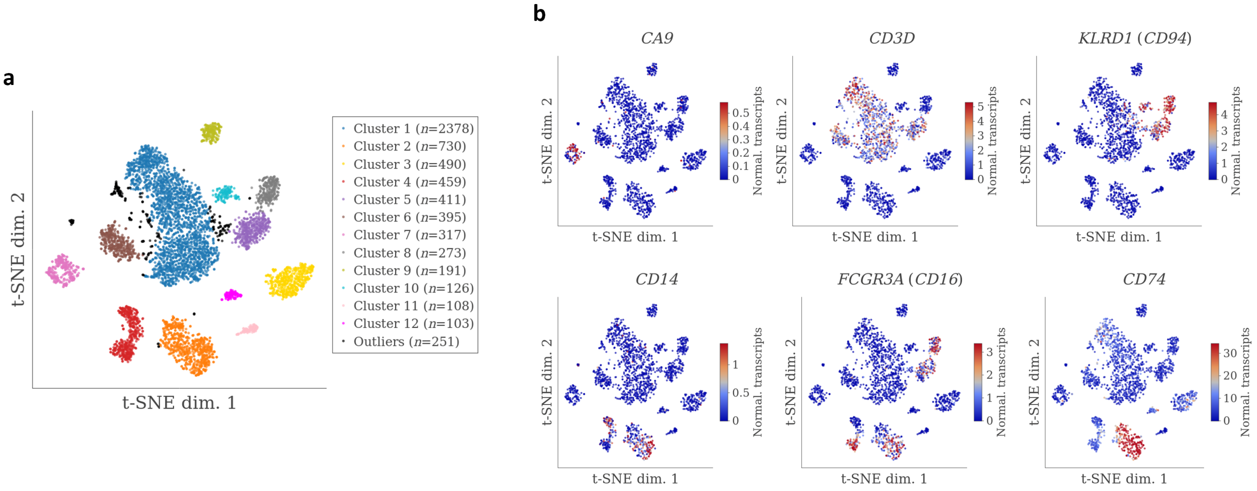


**Supplementary Figure** 1: Clustering of kidney cancer data (6,232 cells) and marker-based cell type identification. **a** t-SNE visualization and DBSCAN clustering results. DBSCAN was applied directly on the t-SNE output with the parameters eps=5.44 and minPts=69. **b** Expression patterns of marker genes, as indicated above each plot, overlaid onto the t-SNE visualization shown in (a). The expression values shown were obtained after applying median normalization to the dataset. For each marker gene, the color scale was clipped at a value corresponding to the mean plus two standard deviations. Only a subset of 2,000 cells, sampled randomly while maintaining cluster proportions, are shown in all plots, to improve readability.

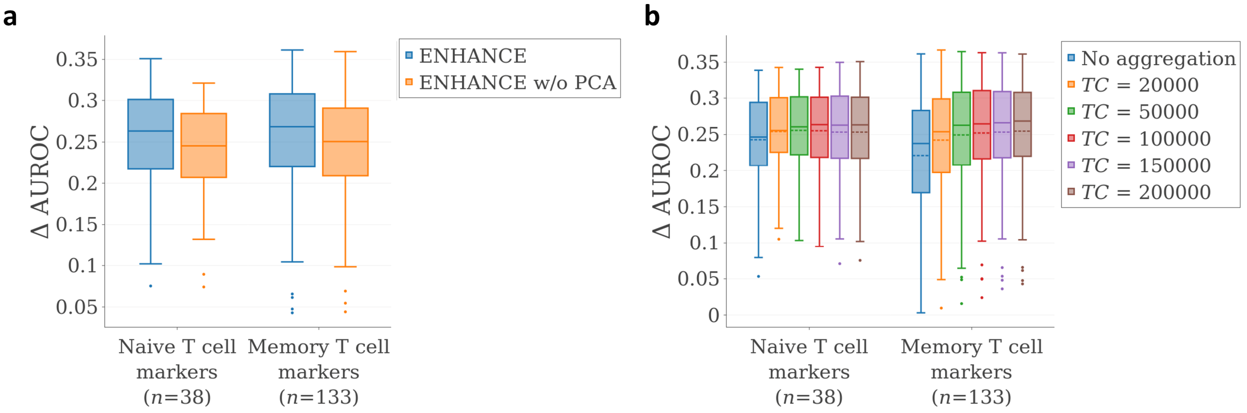


**Supplementary Figure 2: Effect of each step of ENHANCE on denoising performance. a** Improvement in AUROC scores for naïve and memory T cell markers, as in **Fig. 1h**, with and without PC extraction step. **b** Improvement in AUROC scores, as in **Fig. 1h**, with different degrees of nearest-neighbor aggregation, determined by the target transcript count (**Methods**).

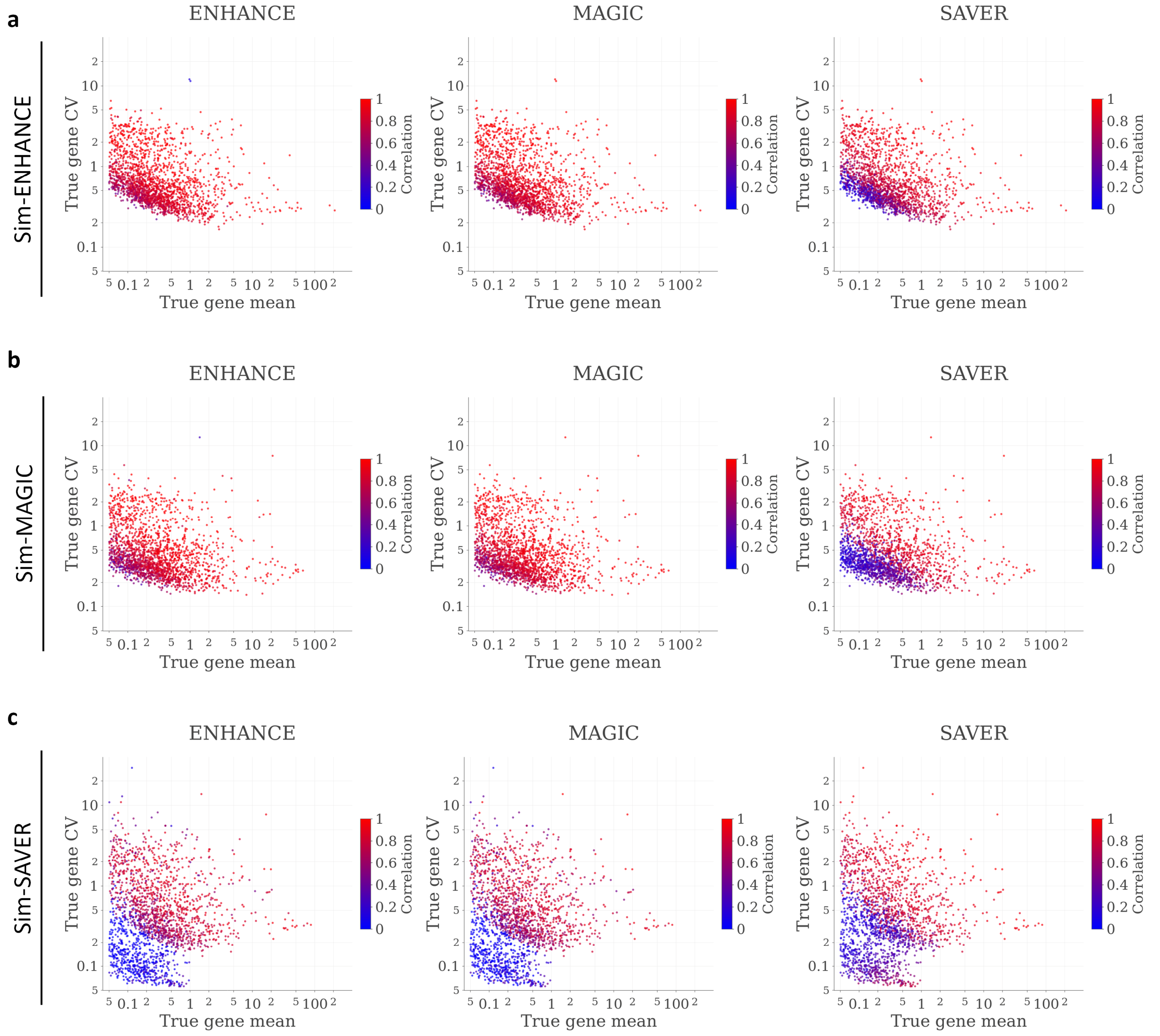


**Supplementary Figure 3: Denoising accuracies on simulated kidney cancer data as a function of gene expression level and variability.** All panels are scatter plots in which dots correspond to expressed genes, and the x- and y-axes correspond to the mean and the coefficient of variation (CV) in the ground truth. Genes are color-coded by Pearson correlation between their gene expression patterns in the ground truth and the denoised data. Results are shown for simulated data generated using Sim-ENHANCE (**a**), Sim-MAGIC (**b**), and Sim-SAVER (**c**). Each column shows results from the denoising method indicated above the plot. For improved readability, only a random subset of 2,000 genes is shown in each plot.

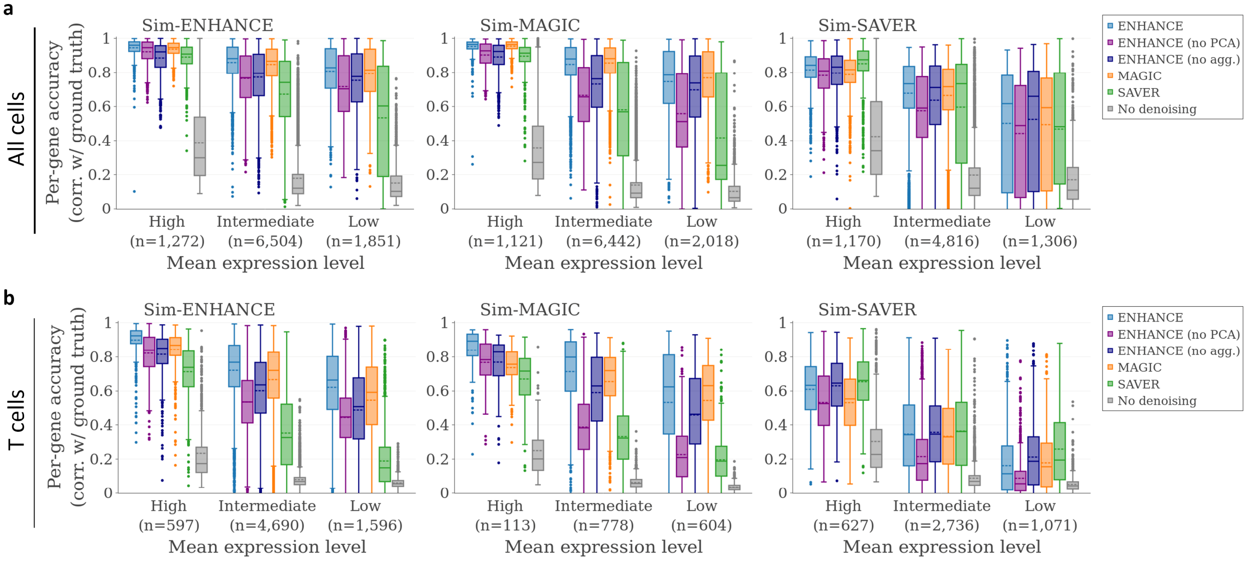


**Supplementary Figure 4: Assessment of denoising accuracies for variants of ENHANCE.**. Shown are gene-wise correlations between ground truth and denoising result, as in **Fig. 2c,d**, obtained using ENHANCE without principal component extraction ("ENHANCE (no PCA)") and ENHANCE without nearest-neighbor aggregation ("ENHANCE (no agg.)"). As a reference, correlations for ENHANCE, MAGIC, SAVER, and no denoising are shown as well. **a** Gene correlations calculated using all cells from the simulated kidney cancer datasets. **b** Gene correlations calculated using only on T cells from the same datasets.

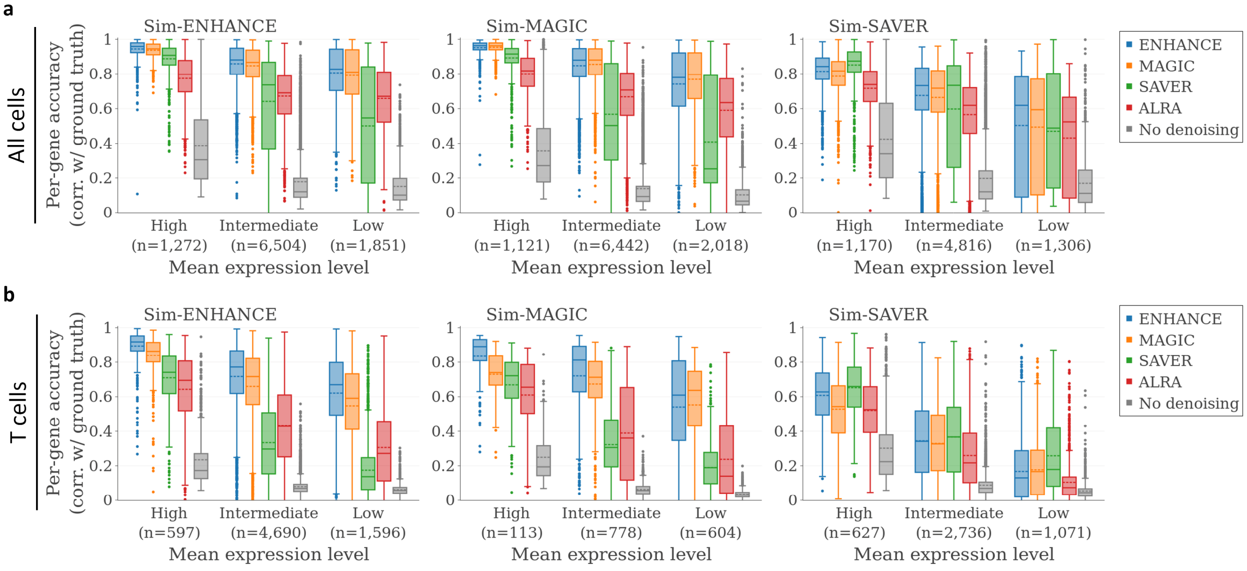


**Supplementary Figure 5: Denoising accuracies on technical replicates of the simulated kidney cancer datasets.** See **Fig. 2c,d** for legend.

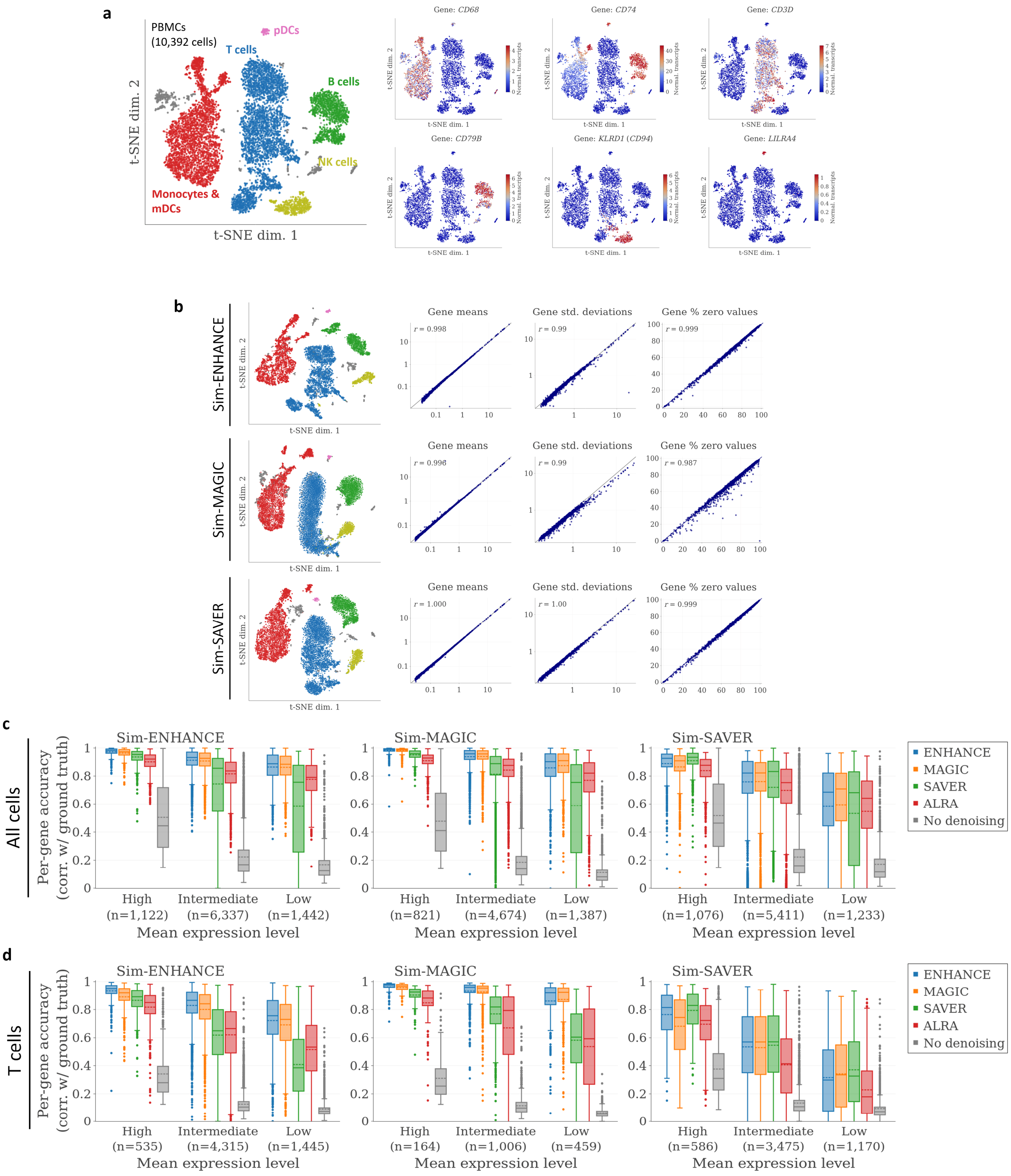


**Supplementary Figure 6: Evaluation of denoising accuracies on simulated PBMC datasets. a** Left: Results of clustering the real PBMC dataset, as in **Fig. 2a**, and cell type annotations. Right: Expression patterns of marker genes used for cell type identification. **b** Left: t-SNE visualizations of datasets simulated using Sim-ENHANCE, Sim-MAGIC, and Sim-SAVER. Right: Comparison of technical characteristics between the simulated and the real data, as in **Fig. 2b**. c Gene correlations between denoised data and ground truth, as in **Fig. 2c**. Gene correlations after selecting T cells, as in **Fig. 2d**.
