## Supplementary Note 1 for "Accurate denoising of single-cell RNA-Seq data using unbiased principal component analysis"

### Supplementary Note 1: Limitations of previously described approaches for simulating scRNA-Seq data in the context of evaluating denoising methods

Huang et al.[1] simulated data by selecting highly expressed genes and cells with large library sizes from the raw data, and treating the raw counts from these genes in these cells as the ground truth. Presumably, the authors reasoned that those measurements were associated with the least amount of technical noise. The authors then simulated sampling noise by drawing from a Poisson distribution (as we do in this work). Additionally, the authors generated artificial efficiency noise by drawing efficiency factors from a Gamma distribution. The resulting datasets only contained data for between 2,059 and 3,529 out of approx. 20,000 protein-coding genes, and for only about 50% of the cells in each real dataset. Therefore, the simulated data only represented limited portions of the biological heterogeneity present in the real data.

Van Dijk et al.[2] simulated data using two different approaches. In the first approach, they obtained microarray expression data representing a time course of developing *C. elegans* worms, and selected genes with dynamic expression patterns. The authors then "downsampled" the expression values "using an exponential distribution such that the result had 80% and 90% of the values set to 0", thus resembling the sparsity of gene expression values observed in single-cell RNA-Seq data. Finally, the data was log-transformed and the expression values of each gene were standardized (transformed to z-scores). Beyond exhibiting a certain proportion of zero values, it is not clear to what extent the resulting data recapitulated the characteristic noise profile of single-cell RNA-Seq data. In the second approach, the authors applied MAGIC to a scRNA-Seq dataset, and treated the resulting dataset as the ground truth (an approach we have adopted in this work). They then again applied "down-sampling using an exponential distribution such that 0%, 60%, 80% and 90% of the values are set to 0", to obtain the simulated data. The authors did not demonstrate that the simulated dataset exhibited similar noise characteristics as the real dataset, for example by comparing the gene standard deviations in the real and the simulated datasets.

Andrews and Hemberg[3] employed three different simulation approaches to examine the effect of denoising methods on differential expression analysis. First, the authors simulated expression matrices containing 1,000 cells and 500 genes by generating synthetic mean values for each gene, introduced a 10-fold expression difference in half of the cells for half of the genes, and simulated transcript counts by drawing from a negative binomial distribution. Second, the authors used Splatter[4] to generate similar matrices that contained 1,000 cells, assigned to 2-10 groups, and 5,000 genes, of which 1-30% had differential expression. When the dropout probability parameter of Splatter is set to zero, it generates transcript counts by drawing from a Poisson distribution, and the authors showed that in this case, the proportion of zero values as a function of Mean expression matches that of real 10x genomics data (Figure S1A in ref.[3]). However, the authors showed that the simulated data did not match real data with respect to other characteristics, such as the distribution of mean expression levels or the mean-variance relationship. Finally, the authors used real datasets from the Tabula Muris consortium and randomly permuted the expression values of genes that did not show evidence of differential expression, in order to test for false positive differential expression results. While the third simulation approach was specifically designed as a negative control, the first two approaches were aimed at simulating realistic scRNA-Seq data. However, neither approach was able to recapitulate the biological heterogeneity of a real scRNA-Seq dataset, as the cell types and their differential expression patterns were completely synthetic. Moreover, the authors did not demonstrate that either approach recapitulated the technical noise characteristics of real scRNA-Seq data, except for the relationship between the mean expression level and the fraction of zero values in one of the simulated datasets from the second approach.

Gong et al.[5] obtained expression matrices after down-sampling the raw sequencing reads. The authors applied this approach to non-UMI-filtered data, to determine the extent to which different denoising methods were able to accurately distinguish between "true" and "technical" zero values. However, for UMI-filtered data, zero values do not have any special significance, and so this approach is inadequate for examining denoising accuracy on UMI-filtered data.

Eraslan et al.[6] employed two different simulation approaches. First, similar to one of the approaches chosen by Andrews and Hemberg[3], the authors used Splatter[4] to generate synthetic expression matrices containing 2,000 cells and 200 genes. Second, the authors adopted the approach described by van Dijk et al. to simulated data based on the C. elegans time-course experiment. We have already discussed the limitations associated with both approaches above.

Talwar et al.[7] simulated datasets by randomly setting between 10-50% of the measurements in real scRNA-Seq datasets to zero. The authors did not demonstrate that this procedure recapitulated the technical noise characteristics of real scRNA-seq data. Based on our current understanding of the sources of technical noise in scRNA-Seq data (**Supplementary Note 1**), this approach cannot be expected to adequately model technical noise in scRNA-Seq data.

Our simulation approach combines individual elements of the previously described approaches: Like Huang et al., but unlike van Dijk et al., we simulate efficiency noise by sampling from a Poisson distribution. Like van Dijk et al., but unlike Huang et al., we use the output of a denoising method as the ground truth. As a result, we are able to simulate datasets that contain data for all genes and cells in the raw data, and we show on a per-gene basis that the resulting data accurately recapitulate the means, standard deviations, and fraction of zero values observed in the raw data.
